# Biocontrol of *Calonectria* causing root rot in *Eucalyptus*

**DOI:** 10.1101/2025.09.02.673731

**Authors:** H. Andrés Villar, Sara García, Elena Tavares, Eduardo Abreo

## Abstract

*Calonectria* spp. are among the most important pathogens affecting Eucalyptus production, causing severe root rot and significant losses in nurseries. In this study, we established a collection of locally isolated strains of *Trichoderma* spp. and *Bacillus* spp., complemented with commercial formulations, to evaluate their potential as biocontrol agents (BCAs) against *Calonectria*. Leaf bating *in vitro* assays were conducted using naturally colonized, non-sterilized substrates to closely simulate nursery conditions. Several locally isolated *Trichoderma* strains (T3, T5, T6, and T7), as well as two commercial products (PC1 and PC2), consistently reduced the incidence of *Calonectria* infection, indicating their capacity to decrease pathogen inoculum levels in the substrate. The reproducibility of these effects across independent trials underscores the robustness of *Trichoderma* as reliable BCAs. Altogether, these findings highlight the potential of locally isolated *Trichoderma* strains as promising alternatives for managing *Calonectria* root rot in Eucalyptus nurseries, while reinforcing the need for further trials under commercial greenhouse conditions to confirm their efficacy in real production systems.

## Introduction

The genus *Eucalyptus* represents a major component of the forestry sector in Uruguay, with *E. grandis* and *E. dunnii* accounting for the largest planted areas in the country (MGAP-DGF, 2025). The cultivation of these species has contributed substantially to the national economy. However, several biotic factors have constrained its productivity, particularly root and stem diseases caused by soilborne pathogens (Tellechea, 2010). Among these, *Calonectria* species are recognized as serious threats to young *Eucalyptus* plants, causing root rot, stem lesions, and damping-off, often resulting in significant plant losses and reduced plantation establishment rates (Alfenas et al., 2015; Bose et al., 2023). This pathogen is one of the most frequently found in Uruguayan forestry nurseries (Martinez et al., 2021).

Management of *Calonectria* in nurseries typically relies on cultural practices and chemical fungicides. However, the use of fungicides is often limited by environmental regulations, concerns over human and environmental health, the development of pathogen resistance and the reduced number of active ingredients, while cultural management is insufficient due to lack of resistant plant genotypes and the pathogen’s survival ability. The utilization of contaminated substrate or soil represents the main source of inoculum (Lui and Chen, 2017), which makes it a critical point in the disease cycle to be targeted. This stage is particularly suitable for control through biological control agents (BCAs) that can suppress pathogen populations.

*Trichoderma* and *Bacillus* species are among the most widely studied BCAs for the management of soilborne plant pathogens, due to their well-documented antagonistic activity and the greater feasibility of formulation for biological control applications (Meher et al., 2020; Liu et al., 2022; Zhang et al., 2023; Yang et al., 2024). Both genera possess multiple mechanisms of action, including mycoparasitism, antibiosis, competition for nutrients and space, and induction of systemic resistance in host plants (Guzman-Guzman et al, 2023; Zhang et al., 2023). Their efficacy has been demonstrated against various pathogens in forestry, horticultural, and agricultural systems, and commercial formulations of both genera are available in many countries. Specifically, various authors have shown that *Calonectria* can be inhibited by the used of *Trichoderma* and *Bacillus* strains (Vitale et al., 2011; Kong and Hong, 2017; Carvalho et al., 2018; Paz et al., 2018; Samavat, 2022; Zang et al., 2023).

The performance of BCAs can be influenced by environmental conditions, substrate and plant characteristics and pathogen populations, making the selection of locally adapted strains essential for effective disease suppression (Hoitink and Boehm 1999; Villar et al., 2022). In this context, the isolation and evaluation of native *Trichoderma* and *Bacillus* strains from *Eucalyptus* nursery environments can increase the likelihood of identifying antagonists with enhanced adaptability and efficacy under local conditions. Including commercially available BCAs in comparative assays provides a performance benchmark and may identify candidates for immediate application in production systems.

The aim of this study was to isolate, identify and evaluate the potential of local *Trichoderma* and *Bacillus* strains to reduce *Calonectria* inoculum under controlled conditions.

## Materials and Methods

### Calonectria

The *Calonectria* isolate 3V from the Biofore Uruguay and INIA Las Brujas-Uruguay collection was used. This isolate was originally obtained from nursery substrate and characterized by its capacity of causing root rot (Villar et al., 2025).

### Trichoderma and Bacillus

A collection of *Trichoderma* and *Bacillus* strains was generated from plant and substrate material obtained from commercial *Eucalyptus* nurseries representing the environments where they are intended to act. The process was carried out in two steps: i) Field sampling. Three-month-old seedlings of *E. dunnii* and *E. grandis* with superior growth and sanitary status were sampled from three commercial *Eucalyptus* nurseries located in the Paysandú region, Uruguay. A total of 125 seedlings were collected (25 for each clone–nursery combination). Additionally, commercial substrate (BCS: 70% composted by pine bark and 30% peat moss; produced by Biofore, Uruguay) widely used in commercial nurseries was also sampled. ii) Isolation. *Trichoderma* and *Bacillus* strains were isolated from the collected seedlings (endophytic -*stem* and *root-* and rhizospheric) and substrate.

For endophyte isolation, stems and roots were washed with tap water, cut into ∼1 cm segments, and surface-sterilized for 120 s in 3% sodium hypochlorite followed by 90 s in 70% ethanol. Samples were rinsed three times in sterile distilled water, and three 100 µL aliquots from the final rinse were plated on potato dextrose agar (PDA; Oxoid, UK) to verify surface sterilization. Stem and root segments were plated on PDA acidified (2 drops of lactic acid per 100 mL) for *Trichoderma* isolation or Luria–Bertani (LB) agar for *Bacillus* isolation, with five stem and five root segments per plate. A total of 25 plates were prepared and incubated at 25 °C.

For isolation from rhizosphere and substrate, the root of each seedling was gently shaken to remove loosely adhering soil, while retaining the soil firmly attached to the roots. Root samples (2.5 g, including adhering soil) were transferred to Erlenmeyer flasks containing 100 mL of sterile saline solution. For substrate-derived isolates, 10.0 g of the BCS were subsampled and suspended in 90 mL of sterile saline solution. All suspensions were agitated at 180 rpm for 20 min. Serial dilutions were prepared up to 10^−3^, and 100 µL aliquots of each dilution were plated in triplicate onto acidified PDA for *Trichoderma* isolation, and LB agar (with thermal shock of 80°C for 30 min) for *Bacillus*.

Plates were incubated at 25 °C and examined daily starting 48 h after plating. Colonies with morphological characteristics consistent with *Trichoderma* or *Bacillus* were subcultured to obtain pure isolates, which were confirmed to genus level by microscopic observation of conidiophores/conidia (*Trichoderma*) or cell/spore morphology (*Bacillus*).

Additionally to the newly isolated strains, three commercially available biological fungicides containing *Trichoderma* or *Bacillus* were included in the evaluation.

### Inoculum production

#### Trichoderma

Inoculum was produced by culturing the strains on PDA plates. Strains were grown in 90 mm Petri dishes at 25 °C. After 7 days, fungal conidia were harvested by scraping the agar surface, suspended in sterile distilled water, and quantified by conidial counting in a Neubauer chamber.

#### Bacillus

Inoculum was produced via submerged fermentation. Colonies from 2-day-old LB agar plates were transferred into Erlenmeyer flasks containing 2XSG broth (composition) and incubated for 72 h at 28 °C with shaking. Serial dilutions (10^?6^, 10^?7^, 10^?8^) were plated on LB agar in triplicate, incubated at 28 °C for 48 h, and colony-forming units (CFU) were enumerated.

#### Calonectria

*Calonectria* 3V was grown in 90 mm Petri dishes with Malt Extract culture medium (Oxoid, UK) at 25 °C for 7 days, after which the agar was cut into small pieces, suspended in sterile distilled water, and blended for 2 min in a previously sterilized blender (Blanco and Romero, 2017). Serial dilutions were performed to confirm a target concentration of ∼1 × 104 CFU/mL.

### Biocontrol assays

Biocontrol efficacy was assessed using the bait plant technique described by Gonçalves et al. (2001), an effective method for quantifying *Cylindrocladium* in soils with varying inoculum densities. Treatments consisted of the co-inoculation of *Trichoderma* or *Bacillus* strains (individually) with the *Calonectria* isolate in plastic trays (13 × 17 cm) containing 200 g of BCS commercial nursery substrate.

*Trichoderma* suspensions were standardized to final concentrations of 5 × 10^7^ conidia/mL (*Trichoderma*) or 5 × 10^7^ CFU/mL (*Bacillus*). Commercial products were adjusted to the same concentrations prior to use. Inoculations with the *Calonectria* strain were performed using suspensions adjusted to ∼1 × 10^4^ CFU/mL.

Each biocontrol tray received 5.0 mL of antagonist suspension and 5.0 mL of pathogen suspension at the concentrations described above, plus 80 mL of sterile water to achieve usual substrate humidity levels. Trays were incubated at 25 °C for 5 days (antagonist–pathogen interaction period). After 5 days, 30 leaf segments of *E. globulus* (∼1.5 × 0.5 cm, previously disinfected by 5 min in 3% sodium hypochlorite) were partially buried in the substrate and trays were re-incubated at 25 °C to allow pathogen infection. From the fourth day after leaf placement, the number of infected leaves per plastic trays was recorded. Control treatments consisted of a disease control with pathogen inoculation only (Only P -5 mL, 1 × 10^4^ CFU/mL-) and a healthy control without pathogen inoculation.

Disease assessment was conducted by a single evaluator on the same day for all treatments, which limited the number of treatments per trial to a maximum of 11. Consequently, four independent biocontrol assays were conducted using the same experimental technique but evaluating different sets of strains in each trial. In trials 1 and 2, *Trichoderma* strains were assessed, while in trials 3 and 4, *Bacillus* strains were tested. In total, 10 *Trichoderma* and 12 *Bacillus* strains were evaluated. From the second trial onward, two additional reference controls were included to estimate pathogen inoculum reduction: Reference 1 (Only P –30%), with 30% less inoculum than the control Only P, and Reference 2 (Only P –70%), with 70% less inoculum than the control Only P was applied. Strain T3 was systematically included in all experiments as a bridging strain to enable cross-trial comparisons. All treatments, both those tested in a single trial and those systematically included across trials (e.g., T3 and reference controls), were replicated five times. For each trail, a different batch of substrate was used and not sterilized to better represent nursery production conditions. Substrate was microbiologically characterized using a representative sample, by quantifying total culturable fungi (in acidified PDA) and bacteria and actinomycetes (in R2A; Oxoid, UK) through the same dilution–plating technique used for substrate-derived isolates described above. From this analysis, bacteria, actinomycetes, and fungi were recovered. Bacteria represented the most abundant group (10^6^ CFU g^−1^), followed by actinomycetes (10^5^ CFU g^−1^) and fungi (10^4^ CFU g^−1^) (Table 1).

**Table 1.**
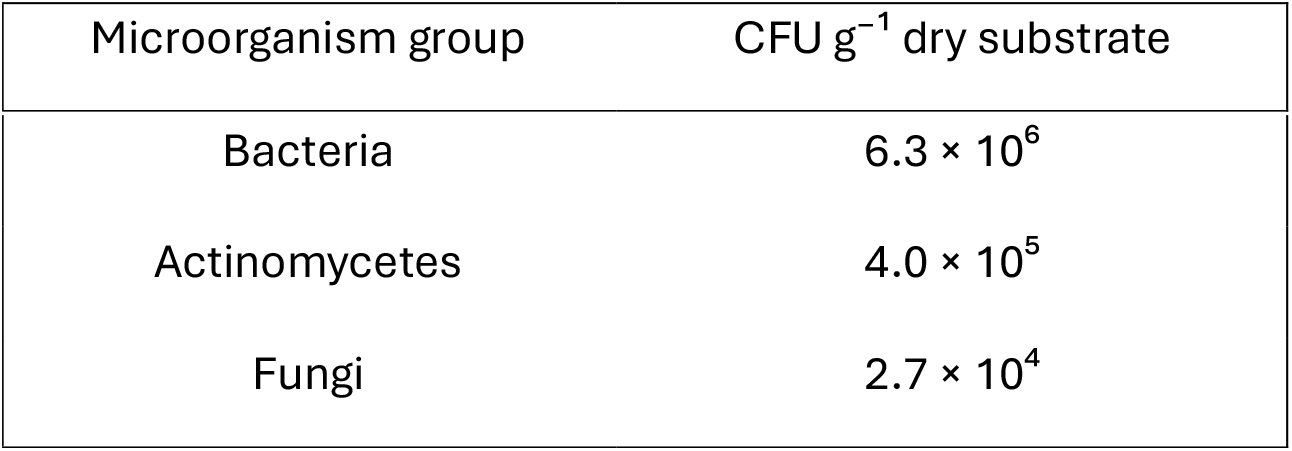
Microbiological characterization of the commercial substrate used in biocontrol assays.

| Microorganism group | CFU g<sup>-1</sup> dry substrate |
| --- | --- |
| Bacteria | $6.3 \times 10^6$ |
| Actinomycetes | $4.0 \times 10^5$ |
| Fungi | $2.7 \times 10^4$ |

### Trichoderma identification

Three strains that showed the greatest antagonistic capacity were subjected to molecular identification by amplification and sequencing of translation elongation factor 1-a (TEF1). Genomic DNA was extracted using the CTAB method described by Murray and Thompson (1980). Trichoderma strains were cultured on PDA medium at 25 °C for 4 days. Mycelium was then harvested with a sterile scalpel, transferred into 1.5 ml Eppendorf tubes, and homogenized using sterile plastic pestles prior to the chemical DNA extraction steps. Amplification of the translation elongation factor 1-a (TEF1) was performed following the protocol described by Kubicek et al. (2003). The PCR was prepared in 50 µl of total volume, 0.25 µl of 5U Dream Taq polymerase (Thermo Fisher Scientific), 5 µl of 2.0 mM of each dNTP solution (Thermo Fisher Scientific), 5 µl of amplification buffer (Thermo Fisher Scientific), and 1.0 µl of DNA (approximately 50 ng/µL), 2.5 µl of each primer EF1-T (5′-ATGGGTAAGGATGACAAGAC-3′) and EF2 (5′-GGA(G/A)GTACCAGT(G/C)ATCATGTT-3′). PCR reaction was run with an annealing temperature of 55 °C. The PCR products were analyzed on agarose gel 1% yielding an amplicon of about ≈900bp.

The amplification products were sequenced by Macrogen Inc. (Seoul, Korea). Sequence similarity searches were performed with BLAST network service in NCBI GenBank (https://blast.ncbi.nlm.nih.gov/Blast.cgi?PROGRAM=blastn&PAGE_TYPE=BlastSearch&LINK_LOC=blasthome). Ex-type cultures sequences (Bissett et al. 2015) and representative sequences of the closest matching species from Genbank were selected to be included in the phylogenetic analysis (Table 2). The sequence alignment was done using Clustal W algorithm within the free software MEGA 12 (Tamura et al. 2021). Manual adjustments were made to improve the alignments. Maximum parsimony (MP) and maximum likelihood (ML) phylogenetic analyses were implemented in MEGA 12. For MP analysis, we used the addition of 1000 random replicates and tree bisection and reconnection (TBR) were selected as branch swap ping algorithm. Gaps were treated as missing character, and all characters were unordered and of equal weight. Branches of zero length were collapsed. To estimate branch support, MP bootstrap values were determined using 1000 bootstrap replicates (Felsenstein 1985). For the ML analysis, the evolutionary history was inferred by using the method based on the Kimura 2-parameter model, with 1000 replicates of bootstrap test. MP and ML analyses generated nearly identical topologies for each data set; thus, only the topology from the ML analysis is presented in the main text.

**Table 2:**
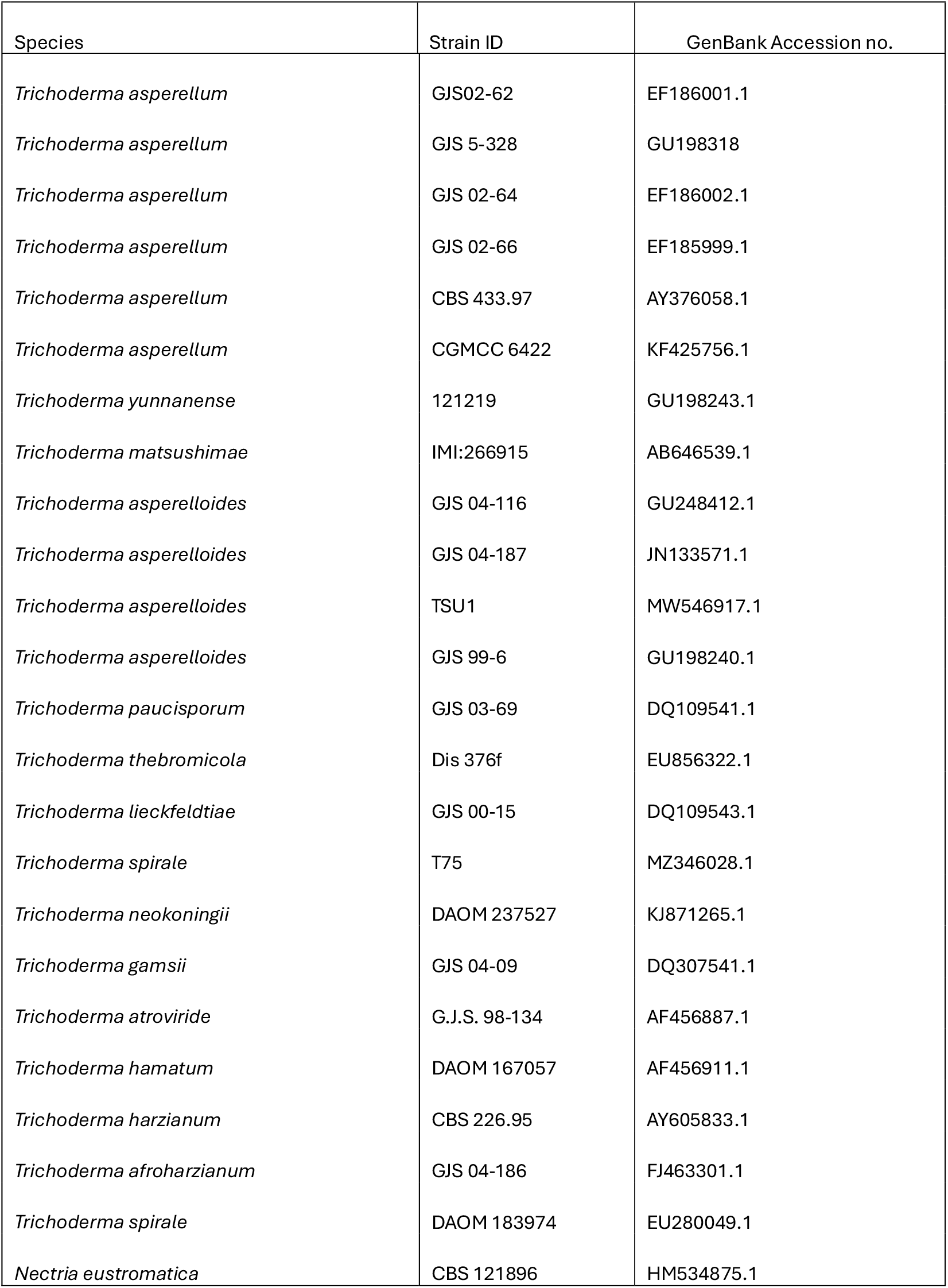
Name and Genbank accession number for the transcription elongation factor (TEF) of reference strains of *Trichoderma* species used in phylogenetic analysis.

### Statistical analysis

The incidence of infection (proportion of infected leaves and non-infected leaves) was analysed using pairwise proportion comparisons (*pairwise.prop.test*) based on the Chi-squared test, with Holm correction for multiple comparisons. For each evaluation day, *p*-value matrices were generated and grouping letters were assigned to treatments according to their statistical diferences (*p* ≤ 0.05). All analyses were performed in R software (version 4.4.2, R Core Team, 2024).

## Results

### *Trichoderma* and *Bacillus* collection

A collection of 20 *Trichoderma* and 46 *Bacillus* strains was obtained from surveys conducted in commercial *Eucalyptus* nurseries. From this collection, strains isolated from different sources (endophytes from stem or root, rhizosphere, and commercial substrate), as well as from different *Eucalyptus* species, clones, and nurseries, were selected for biocontrol assays to capture the widest possible diversity of origins for both genera (Table 1). In addition, commercially available products registered in Uruguay as biological fungicides based on *Trichoderma* (PC1, PC2) and *Bacillus* (PC3) strains were included (Table 1).

**Table 1.** Strains of *Trichoderma* and *Bacillus* selected for the four biocontrol assays.

| Trial | Strain ID | Source | *Eucalyptus* | Nursery |
| --- | --- | --- | --- | --- |
| 1 | T12 | Root | *E. grandis* | B |
|  | T13 | Root | *E. dunnii* – | C |
|  | T3 | Stem | *E. dunnii* – | B |
|  | T4 | Rhizosphere | *E. grandis* | B |
|  | T5 | Rhizosphere | *E. dunnii* – | C |
| 2 | T7 | Rhizosphere | *E. dunnii* – | B |
|  | T6 | Rhizosphere | *E. dunnii* – | C |
|  | T16 | Substrate | – | C |
|  | PC1 | – | – | – |
|  | PC2 | – | – | – |
| 3 | B1 | Stem | *E. dunnii* – | B |
|  | B29 | Root | *E. dunnii* – | A |
|  | B5 | Root | *E. dunnii* – | B |
|  | B15 | Rhizosphere | *E. dunnii* – | A |
|  | B19 | Rhizosphere | *E. dunnii* – | A |
|  | B39 | Stem | *E. dunnii* – | B |
| 4 | B35 | Root | *E. dunnii* – | C |
|  | B36 | Root | *E. dunnii* – | C |
|  | B42 | Stem | *E. dunnii* – | A |
|  | B43 | Stem | *E. grandis* | B |
|  | B23 | Rhizosphere | *E. dunnii* – | A |
|  | PC3 | – | – | – |

### Biocontrol

Among the selected isolates, several *Trichoderma* strains reduced the percentage of pathogen-infected leaves (Figure 1). In Trial 1, strains T3 and T5 showed significantly lower infection levels compared to the control. In Trial 2, all *Trichoderma* strains tested decreased infection incidence caused by *Calonectria*, with T6, T7, PC2, and PC1 achieving the strongest reductions. The infection levels recorded for these strains were comparable to those obtained with T3 and with the treatment where the pathogen was inoculated at a 70% lower concentration than the control (Only P -70%).

**Figure 1.**
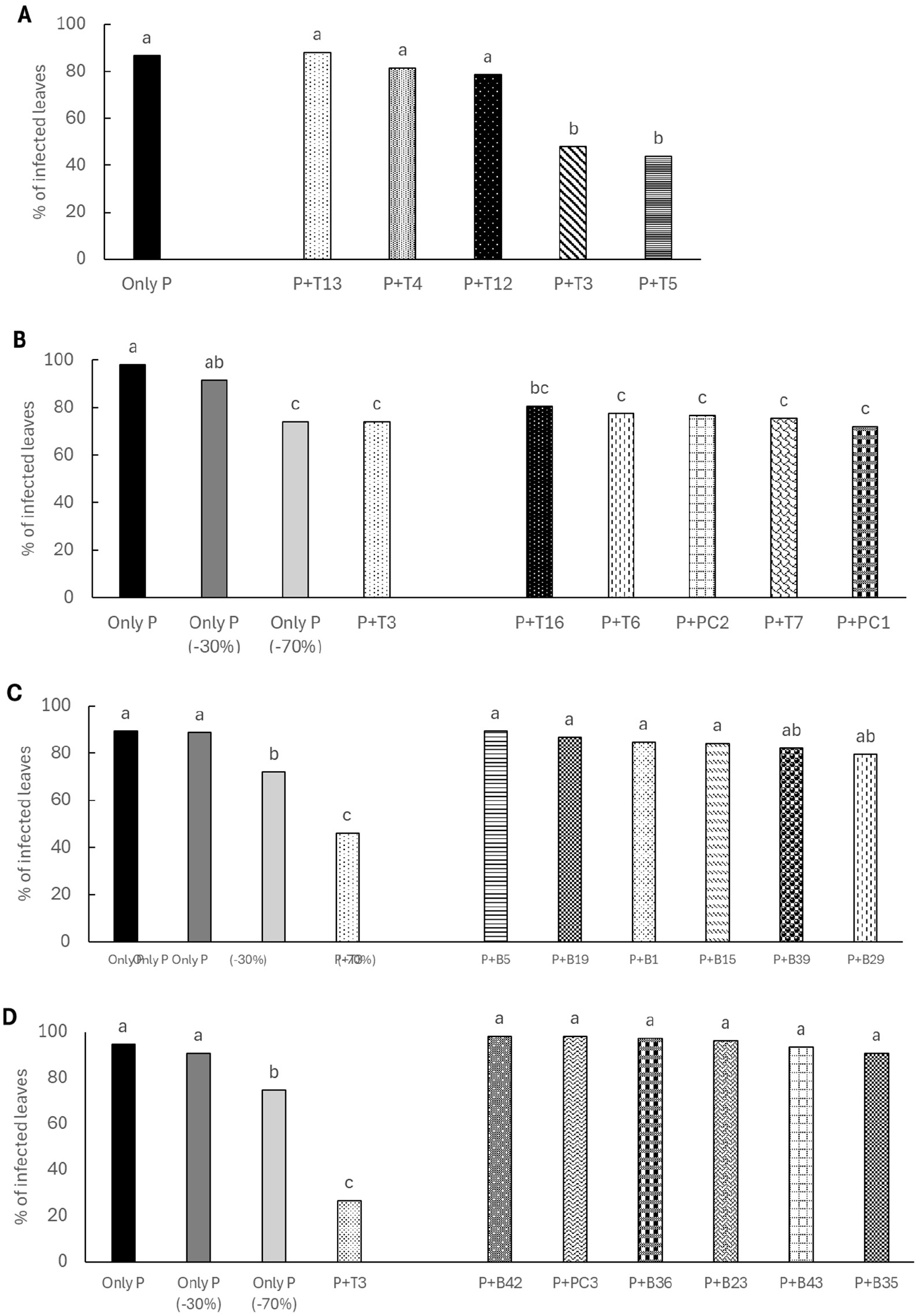
Percentage of infected leaves the in four independent trials according to *Trichoderma* or *Bacillus* strains. In trials 1 (A) and 2 (B) were evaluated *Trichoderma* strains, and trials 3 (C) and 4 (D) *Bacillus* strains. Bars represent the mean percentage of infected leaves for each treatment. Different letters above the bars indicate statistically significant differences between treatments (p < 0.05).

In contrast, none of the *Bacillus* strains were able to reduce the incidence of infected leaves. Notably, in all cases where strain T3 (used as a bridging strain) was inoculated, the percentage of infected leaves was significantly lower than the diseased control (Only P), and equal to or lower than that observed in the Only P – 70% treatment.

### Trichoderma identification

The TEF sequences analysis involved three of local strains of *Trichoderma* that showed the greatest antagonistic capacity (T3, T5 and T7), 23 *Trichoderma* sequences from GeneBank, and *Nectria eustromatica* as out group. The analysis involved 26 nucleotide sequences. Codon positions included were 1st + 2nd + 3rd + Noncoding. All positions containing gaps and missing data were eliminated. There were a total of 490 positions in the final dataset. The phylogenetic tree showed that the 3 isolates belonged to two species of *Trichoderma* (Figure 2). Strains T3 and T5 formed a clade with *Trichoderma asperellum*, and T7 grouped with *Trichoderma asperelloides.*

**Figure 2.**
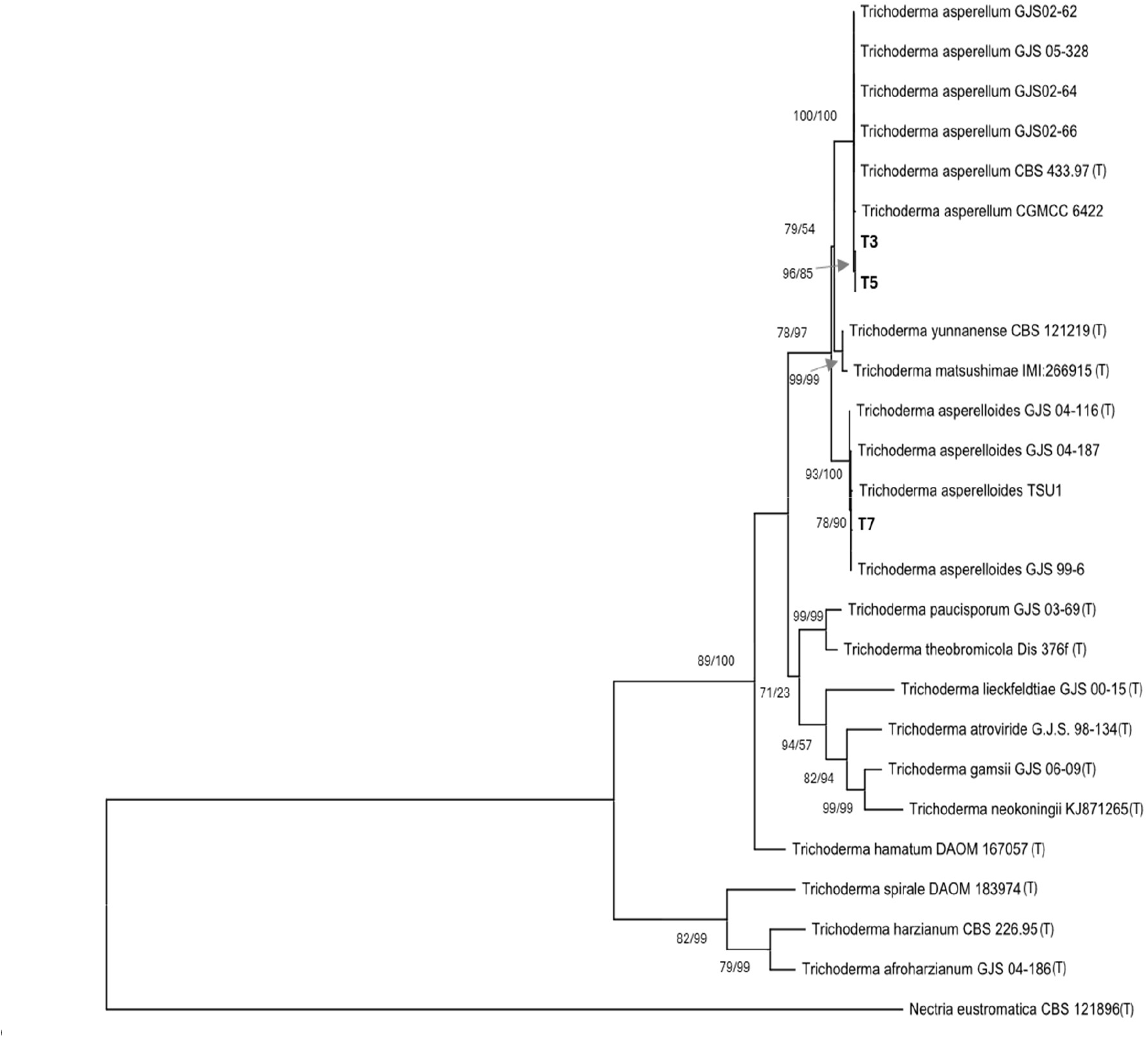
Maximum likelihood tree based on partial sequences of the translation elongation factor 1 alfa (tef1alfa) of local sequences and sequences retrieved from genBank including ex-type strain sequences (T). Bootstrap values of 1000 replicates of the ML/MP analysis are shown below branches. only values greater than 70 are presented. *Nectria eustromatica* was used as outgroup.

## Discussion

The search for local microbial control agents is justified since sustainable alternatives to chemical active ingredients demands the selection of adapted microbial strains, with higher chances of effective control in productive environments. In this work, we set up a collection of locally isolated *Trichoderma* spp. and *Bacillus* spp. strains obtained from different sources, to which commercially available formulations were added for comparison. Among these, both locally isolated *Trichoderma* strains (T3, T5, T6, and T7) and commercial products (PC1 and PC2) consistently achieved significant reductions in *Calonectria* infection, indicating their ability to decrease pathogen inoculum levels in the substrate.

These results are consistent with previous reports highlighting *Trichoderma* spp. as promising antagonists of soilborne pathogens (Harman, 2005; Muhammad et al., 2018). Several studies have reported *Trichoderma* strains as antagonists of *Calonectria* (Vitale et al., 2011; Kong and Hong, 2017; Carvalho et al., 2018). Interestingly, strain T3 consistently reduced the incidence of infected leaves across all trials in which it was included, demonstrating the reproducibility of its biocontrol effect. Although some variation was observed in its efficacy, infection levels were always equal to or lower than those obtained with the disease control treatment in which the pathogen inoculum was reduced by 70% (Only P –70% treatment), which could be taken as an indication of inoculum reduction by at least 70% when T3 was applied. Variability observed around 70% reduction may be attributed to differences among substrate batches used in each trial. Even though commercial substrates are standardized, factors such as storage time, temperature, and humidity can lead to diferences in their microbiological, chemical, and physical properties, potentially influencing pathogen development and biocontrol performance. Further studies are required to identify the specific conditions that enable this strain to fully express its antagonistic potential.

Moreover, other strains performed at levels comparable to the Only P (–70%) control suggests that they may achieve medium-to-high reductions (>70%) in pathogen inoculum. Importantly, these performances were achieved in a non-sterilized substrate, meaning that the biocontrol strains not only antagonized the pathogen but also successfully competed with the naturally occurring microbial community. According to the substrate sample evaluated, this community was abundant, with total culturable bacteria reaching 10^6^, actinomycetes 10^5^, fungi 10^4^ CFU g^−1^. These findings strengthen the case for their application as promising and effective management alternatives under nursery production conditions.

The inclusion of commercial products based on *Trichoderma* with biocontrol capacity brings the adoption of biological control in *Eucalyptus* nurseries closer to reality, since it would not require going through the entire process of product development and registration. However, this does not preclude the use of locally isolated strains identified in this study, but rather broadens the spectrum of available microbial resources, with strains from different Trichoderma species. The coexistence of commercial formulations and newly isolated strains could provide complementary tools, strengthening the foundation for the gradual incorporation of biological control into nursery disease management programs in Uruguay.

In contrast, none of the *Bacillus* strains tested here significantly reduced the number of leaves infected, even though *Bacillus* spp. are widely recognized as effective biocontrol agents, also against *Calonectria* (Paz et al., 2018; Samavat, 2022; Zang et al., 2023). The absence of activity under our experimental conditions should not be interpreted as definitive lack of potential. Several key factors may have limited their observed efficacy. For instance, the temperature of 25 °C was used because it corresponds to the average conditions registered in the greenhouse. However, the optimal temperature range for *Bacillus* species growth is generally reported as 28–35 °C (Errington and van der Aart, 2020; Ajuna et al., 2020. Additionally, suboptimal formulation or inoculum density, the lack of biofilm-enhancing conditions, low competition ability in the studied substrate, among others, are known to influence colonization and antagonistic activity, may have compromised outcome (Fessia et al., 2022). Therefore, their biocontrol potential cannot be ruled out, and should be re-evaluated under optimized conditions.

Regarding the identification of *T. asperellum* and *T. asperelloides*, these are closely related sister species within the *T. asperellum sensu lato* group (Samuels and Ismaiel, 2010). Their observed biocontrol performance is consistent with the well-established role of *T. asperellum sensu lato* as one of the most widely employed biocontrol species, due to its versatile antagonistic mechanisms that include mycoparasitism, competition for resources, production of antifungal enzymes (e.g., chitinases and glucanases), and induction of host defense responses (Rivera Méndez et al., 2020; Mukherjee, 2022; Abbas et al., 2022). Although no specific reports were found on *T. asperellum* or *T. asperelloides* directly antagonizing *Calonectria*, both species have been repeatedly associated with antagonism against other soilborne pathogens in diverse plant–pathogen systems (Phoka et al., 2020; Rivera Méndez et al., 2020; Boukaew et al., 2024; Hasna et al., 2025), highlighting their strong antifungal potential against this type of pathogens. To our knowledge, this is the first report of strains belonging to *T. asperellum* and *T. asperelloides* exhibiting antagonistic activity against *Calonectria* pathogenic species of *Eucalyptus.*In summary, this study identified several *Trichoderma* strains with promising antagonistic capacity. One of them, *T. asperellum* strain T3 isolated from *Eucalyptus* stems performed consistently across independent trials. These results were obtained under *in vitro* conditions using a method that goes beyond traditional dual culture assays, since it quantified pathogen inoculum and was carried out in a non-sterilized commercial substrate with a natural microbiota, under temperatures representative of greenhouse conditions. This approach brings the evaluation closer to real production scenarios, thereby strengthening the evidence of their potential efficacy in nurseries. Altogether, these findings suggest that the selected strains are likely to maintain consistent performance under nursery conditions. Nonetheless, greenhouse trials remain necessary to confirm their effectiveness under real production conditions.

## Acknowledgments

We gratefully acknowledge UPM Biofore and the National Institute of Agriculture Research (INIA) for funding this project, as well as the staff members of both institutions involved in its execution.

## References

Abbas, A., Mubeen, M., Zheng, H., Sohail, M. A., Shakeel, Q., Solanki, M. K., Iftikhar, Y., Sharma, S., Kashyap, B. K., Hussain, S., Zuñiga Romano, M.C., Moya-Elizondo, E. A., & Zhou, L. (2022). Trichoderma spp. genes involved in the biocontrol activity against Rhizoctonia solani. Frontiers in Microbiology, 13, 884469. 10.3389/fmicb.2022.884469

Ajuna, H. B., Kim, I., Han, Y. S., Maung, C. E. H., & Kim, K. Y. (2020). Aphicidal activity of Bacillus thuringiensis strain AH-2 against cotton aphid (Aphis gossypii). Entomological Research, 50(11), 559–566. 10.1111/1748-5967.12481

Alfenas, A. C., Lombard, L., Pereira, O. L., Alfenas, R. F., & Crous, P. W. (2015). Species of Calonectria associated with Eucalyptus leaf blight and root rot in Brazil. Persoonia, 35, 127–146. 10.3767/003158515X687669

Blanco Vieira, M. (2017). Evaluación de la patogenicidad de aislados de Calonectria spp. en Eucalyptus spp. en Uruguay (Tesis de Maestría). Universidad de la República, Montevideo, Uruguay.

Bissett, J., Gams, W., Jaklitsch, W., & Samuels, G. J. (2015). Accepted Trichoderma names in the year 2015. IMA Fungus, 6(2), 263–295. 10.5598/imafungus.2015.06.02.02

Bose, T., Roy, S., & Sharma, P. (2023). Genetic diversity and pathogenic potential of Calonectria species associated with Eucalyptus. Annals of Applied Biology, 183(3), 1–13. 10.1111/aab.12800

Boukaew, S., Chumkaew, K., Petlamul, W., Srinuanpan, S., Nooprom, K., & Zhang, Z. (2024). Biocontrol effectiveness of Trichoderma asperelloides SKRU-01 and T. asperellum NST-009 on postharvest anthracnose in chili pepper. Food Control, 163, 110490. 10.1016/j.foodcont.2024.110490

Cai, F., & Druzhinina, I. S. (2021). In honor of John Bissett: Authoritative guidelines for the molecular identification of Trichoderma. Fungal Diversity, 107, 1–69. 10.1007/s13225-021-00475-y

Carvalho, D. D. C., Oliveira, D. F., de Souza, R. M., da Silva, L. R., & Campos, V. P. (2018). Biological control of leaf spot and growth promotion of Eucalyptus plants by Trichoderma spp. Forest Pathology, 48(2), e12397. 10.1111/efp.12397

Errington, J., & van der Aart, L. T. (2020). Microbe Profile: Bacillus subtilis: Model organism for cellular development, and industrial workhorse. Microbiology, 166(5), 425–427. 10.1099/mic.0.000922

Felsenstein, J. (1985). Confidence limits on phylogenies: An approach using the bootstrap. Evolution, 39(4), 783–791. 10.1111/j.1558-5646.1985.tb00420.x

Fessia, A., Sartori, M., García, D., Fernández, L., Ponzio, R., Barros, G., & Nesci, A. (2022). In vitro studies of biofilm-forming Bacillus strains, biocontrol agents isolated from the maize phyllosphere. Biofilm, 4, 100097. 10.1016/j.bioflm.2022.100097

Gonçalves, R. C., Alfenas, A. C., Maffia, L. A., & Crous, P. W. (2001). Evaluation of bioassays to quantify Cylindrocladium inocula in soil. Mycoscience, 42(3), 261–264. 10.1007/BF02463986

Guzmán-Guzmán, P., Kumar, A., de los Santos-Villalobos, S., Parra-Cota, F. I., Orozco-Mosqueda, M. C., Fadiji, A. E., Hyder, S., Babalola, O. O., & Santoyo, G. (2023). Trichoderma species: Our best fungal allies in the biocontrol of plant diseases—A review. Plants, 12(3), 432. 10.3390/plants12030432

Harman, G. E. (2005). Overview of mechanisms and uses of Trichoderma. Phytopathology, 96(2), 190–194. 10.1094/PHYTO-96-0190

Hasna, Z., Fathia, A., et al. (2025). Biocontrol efficacy of Trichoderma asperellum against Fusarium wilt in tomato: Induction of resistance mechanisms and growth promotion. Discover Plants, 2, 136. 10.1007/s44372-025-00224-1

Hoitink, H. A. J., & Boehm, M. J. (1999). Biocontrol within the context of soil microbial communities: A substrate-dependent phenomenon. Annual Review of Phytopathology, 37, 427–446. 10.1146/annurev.phyto.37.1.427

Kong, P., & Hong, C. (2017). Biocontrol of boxwood blight by Trichoderma koningiopsis. Plant Disease, 101(6), 858–863. 10.1094/PDIS-10-16-1476-RE

Kubicek, C. P., Bissett, J., Druzhinina, I., Kullnig-Gradinger, C. M., & Szakacs, G. (2003). Genetic and metabolic diversity of Trichoderma: A case study on south-east Asian isolates. Fungal Genetics and Biology, 38(3), 310–319. 10.1016/S1087-1845(02)00583-2

Li, S., Zhang, F.-M., Shang, X.-J., & Hou, R. (2023). Control effect and mechanism of Trichoderma asperellum TM11 against blueberry root rot. Polish Journal of Microbiology, 72(3), 325–337. 10.33073/pjm-2023-034

Liu, Q.-L., & Chen, S.-F. (2017). Two novel species of Calonectria isolated from soil in a natural forest in China. MycoKeys, 26, 25–60. 10.3897/mycokeys.26.14688

Liu, Y., He, P., Munir, S., Ahmed, A., Wu, Y., Yang, Y., Lu, J., Wang, J., Yang, J., Pan, X., Tian, Y., & He, Y. (2022). Potential biocontrol efficiency of Trichoderma species against oomycete pathogens. Frontiers in Microbiology, 13, 974024. 10.3389/fmicb.2022.974024

Martínez, G., Escudero, P., Balero, R., Croci, R., Lupo, S., & Martínez Haedo, J. (2021). La sanidad en viveros forestales: un caso exitoso de construcción interinstitucional. Revista INIA Forestal, 67, 20–28.

Meher, J., Rajput, R. S., Bajpai, R., Teli, B., & Dashora, K. (2020). Trichoderma: A globally dominant commercial biofungicide for sustainable agriculture. In Recent Advances in Applied Microbiology (pp. 199–218). Springer. 10.1007/978-3-030-54758-5_9

MGAP-DGF. (2025). Superficie forestal del Uruguay (Bosques Plantados): Período 1975–2023. Ministerio de Ganadería, Agricultura y Pesca, Dirección General Forestal. Montevideo, Uruguay.

Muhammad, S., Abdullah, N. S., Prismantoro, D., Dwisandi, R. F., Safitri, R., Mohd-Yusuf, Y., Mohd Suhaimi, N. S., & Doni, F. (2018). Mechanisms of action and biocontrol potential of Trichoderma against Fusarium in horticultural crops. Biological Control, 123, 18–27. 10.1016/j.biocontrol.2018.04.013

Mukherjee, P. K., Mendoza-Mendoza, A., Zeilinger, S., & Horwitz, B. A. (2022). Mycoparasitism as a mechanism of Trichoderma-mediated suppression of plant diseases. Fungal Biology Reviews, 39, 15–33. 10.1016/j.fbr.2021.11.004

Murray, M. G., & Thompson, W. F. (1980). Rapid isolation of high molecular weight plant DNA. Nucleic Acids Research, 8(19), 4321–4325. 10.1093/nar/8.19.4321

Paz, I. C. P., Santin, R. C. M., Guimarães, A. M., Rosa, O. P. P., Quecine, M. C., Silva, M.C.P.E., Azevedo, J. L., & Matsumura, A. T. S. (2018). Biocontrol of Botrytis cinerea and Calonectria gracilis by eucalyptus growth promoters Bacillus spp. Microbial Pathogenesis, 121, 106–109. 10.1016/j.micpath.2018.05.026

Phoka, N., Suwannarach, N., Lumyong, S., Ito, S., Matsui, K., Arikit, S., & Sunpapao, A. (2020). Role of volatiles from the endophytic fungus Trichoderma asperelloides PSU-P1 in biocontrol potential and in promoting the plant growth of Arabidopsis thaliana. Journal of Fungi, 6(4), 341. 10.3390/jof6040341

Qin, W.-T., & Zhuang, W.-Y. (2016). Seven wood-inhabiting new species of the genus Trichoderma in the Viride clade. Scientific Reports, 6, 27074. 10.1038/srep27074

Rivera-Méndez, W., Obregón, M., Morán-Diez, M. E., Hermosa, R., & Monte, E. (2020). Trichoderma asperellum biocontrol activity and induction of systemic defenses against Sclerotium cepivorum in onion plants under tropical climate conditions. Biological Control, 141, 104145.10.1016/j.biocontrol.2019.104145

Samavat, S. (2022). Antagonistic activity of Bacillus isolates against Calonectria pseudonaviculata, the causal agent of boxwood blight. Iranian Journal of Forest and Poplar Research, 30(2), 164–175.

Samuels, G. J., & Ismaiel, A. (2010). Trichoderma asperellum sensu lato consists of two cryptic species. Mycologia, 102(4), 944–966. 10.3852/09-243

Santoyo, G., Orozco-Mosqueda, M. C., Afridi, M. S., Mitra, D., Valencia-Cantero, E., & Macías-Rodríguez, L. (2024). Trichoderma and Bacillus multifunctional allies for plant growth and health in saline soils: Recent advances and future challenges. Frontiers in Microbiology, 15, 1423980. 10.3389/fmicb.2024.1423980

Tamura, K., Stecher, G., & Kumar, S. (2021). MEGA11: Molecular evolutionary genetics analysis version 11. Molecular Biology and Evolution, 38(7), 3022–3027. 10.1093/molbev/msab120

Tellechea, N. (2010). Algunas enfermedades y plagas de Eucalyptus y Pinus en Uruguay. En XXIV Jornadas Forestales de Entre Ríos. Concordia, Argentina.

Villar, H. A., Vero, S., Pereyra, S., Altier, N., De Lucca, F., Abreo, E., & Pérez, C. A. (2022). Characterization of the antagonistic capacity of Trichoderma spp. from agricultural systems. International Journal of Pest Management, 68(4), 359–368. 10.1080/09670874.2022.2123568

Villar, A., García, S., Centurión, C., Tavares, E., & Abreo, E. (2025, agosto). Biocontrol de cepas de Calonectria causantes de pudrición radicular en Eucalyptus [Resumen en actas de congreso]. Congreso Brasileño de Fitopatología, Lavras, Brasil.

Vitale, A., Cirvilleri, G., Castello, I., Aiello, D., & Polizzi, G. (2012). Evaluation of Trichoderma harzianum strain T22 as a biological control agent of Calonectria pauciramosa. BioControl, 57(5), 687–696. 10.1007/s10526-011-9423-1

Yang, F., Wang, X., Jiang, H., Yao, Q., Liang, S., Chen, W., Shi, G., Tian, B., Hegazy, A., & Ding, S. (2024). Mechanism of a novel Bacillus subtilis JNF2 in suppressing Fusarium oxysporum f. sp. cucumerium and enhancing cucumber growth. Frontiers in Microbiology, 15, 1459906. 10.3389/fmicb.2024.1459906

Zang, Y., Li, X., Zhou, Y., Chen, J., & Zhang, Y. (2023). Biocontrol mechanisms of Bacillus spp. in agriculture. Frontiers in Microbiology, 14, 1173217. 10.3389/fmicb.2023.1173217

Zhang, N., Wang, Z., Shao, J., Xu, Z., Liu, Y., Xun, W., Miao, Y., Shen, Q., & Zhang, R. (2023). Biocontrol mechanisms of Bacillus: Improving the efficiency of green agriculture. Microbial Biotechnology, 16(12), 2250–2263. 10.1111/1751-7915.14348

